# Prioritization of sites for plant species restoration in the Chilean Biodiversity Hotspot: A spatial multi-criteria decision analysis approach

**DOI:** 10.1101/026716

**Authors:** Ignacio C. Fernández, Narkis S. Morales

**Author notes:** **Author contributions**: IF and NM conceived and designed the research; NM performed the niche modeling; IF built the spatial layers and performed the GIS analyses; IF and NM wrote and edited the manuscript.

## Abstract

Various initiatives to identify global priority areas for conservation have been developed over the last 20 years (e.g. Biodiversity Hotspots). However, translating this information to actionable local scales has proven to be a major task, highlighting the necessity of efforts to bridge the global-scale priority areas with local-based conservation actions. Furthermore, as these global priority areas are increasingly threatened by climate change and by the loss and alteration of their natural habitats, developing additional efforts to identify priority areas for restoration activities is becoming an urgent task. In this study we used a Spatial Multi-Criteria Decision Analysis (SMCDA) approach to help optimize the selection of sites for restoration initiatives of two endemic threatened flora species of the “Chilean Winter Rainfall-Valdivian Forest” Hotspot. Our approach takes advantage of freely GIS software, niche modeling tools, and available geospatial databases, in an effort to provide an affordable methodology to bridge global-scale priority areas with local actionable restoration scales. We used a set of weighting scenarios to evaluate the potential effects of short-term *vs* long-term planning perspective in prioritization results. The generated SMCDA was helpful for evaluating, identifying and prioritizing best suitable areas for restoration of the assessed species. The method proved to be simple, transparent, cost effective and flexible enough to be easily replicable on different ecosystems. This approach could be useful for prioritizing regional-scale areas for species restoration in Chile, as well as in other countries with restricted budgets for conservation efforts.

## Implications for Practice

- Developing methodological approaches to identify and prioritize areas for restoration activities is a crucial task for restoration planning, especially in regions with limited resources for conservation initiatives.
- The increasing availability of free GIS software, niche modeling tools, and geospatial databases offer valuable resources that can be integrated into a spatial multi-criteria decision analysis (SMCDA) to help in the selection of best areas for restoration initiatives.
- The SMCDA provide a flexible, transparent, affordable and replicable framework to prioritize regional-scale areas for restoration of plant species in Chile, as well as in other countries with restricted budgets for conservation efforts.

## Introduction

Humans have extensively and profoundly altered Earth’s landscapes by transforming natural habitats in productive lands and by appropriating a vast extension of available natural resources (Ellis et al. 2010; Ellis & Ramankutty 2008). The replacement of natural habitats by agricultural lands, artificial forests, and urban areas has generated the loss, fragmentation, and degradation of natural habitats leading to an alarming global increase in species extinction rates (Foley et al. 2005; Brook et al. 2008). While the current biodiversity crisis is far reaching and complex, resources available for planning and developing conservation strategies are still very limited (Wilson et al. 2009; Watson et al. 2011). Therefore, one of the main challenges for conservation biologist has been the development of methodological approaches that helps in the prioritization of limited resources among different conservation options (Redford et al. 2003).

Among these challenges, a frequent task for conservationists has been the selection of areas to be prioritized for conservation actions. At the global scale these initiatives have aimed to identify those ecoregions that are most valuable for conservation around the world (Brooks et al. 2006). Examples of these initiatives are the “Global 200 Ecoregion” (Olson & Dinerstein 2002), the “Crisis Ecoregion” (Hoekstra et al. 2005), and probably the most recognized of all, the world “Biodiversity Hot Spots” (Myers 1990; Myers et al. 2000; Mittermeier et al. 2004). Even though these initiatives have been fruitful in signaling priority areas at a global-scale, their real success has been criticized because they have not provided information regarding how to allocate resources within prioritized ecoregions (Wilson et al. 2006). For these initiatives to have real impacts in local conservation actions will require the development of complementary regional and local scales prioritization approaches (Redford et al. 2003). This is a critical issue because the large extent of prioritized areas are in developing countries (Brooks et al. 2006) where budgets for conservation initiatives are often much smaller than is required (Waldron et al. 2013).

At finer scales conservationists have placed a great extent of their efforts to identify and bring under protection the most valuable sites for conservation. These efforts have largely been guided by the use of “systematic conservation planning ” (Margules & Pressey 2000), which has provided a useful framework to optimize the selection of sites to be targeted for developing conservation strategies (Sarkar et al. 2006; Wilson et al. 2009). In general conservation planning can be defined as “the process of deciding where, when and how to allocate limited conservation resources to minimize the loss of biodiversity, ecosystem services and other valued aspects of the natural world” (Pressey & Bottrill 2009). It concerns the prioritization of sites based in their biodiversity value, and the participatory planning and collaborative implementation of strategies that secure the long-term viability of biological diversity (Kukkala & Moilanen 2013). Whereas systematic conservation planning has largely influenced the way institutions and governments prioritize the efforts to protect valuable ecosystems (Sarkar et al. 2006), this systematic approach seems to not have permeated to other fundamental conservation actions, such as restoration planning (but see Noss et al. 2009)

Biodiversity restoration activities are among the most expensive conservation strategies worldwide (Holl et al. 2003). However the development of approaches specifically aimed to prioritize sites for restoration or reintroduction of species has been scarcely addressed (Noss et al. 2009). In contrast to the predominant systematic conservation planning approach that focuses primarily in prioritizing areas that currently contain target species, restoration activities often need to prioritize sites that have reduced populations, or even the complete absence of the species to conserve. Furthermore, because systematic conservation planning has focused primarily on current biodiversity patterns, it has had limited applications for conservation strategies in a rapidly changing climate (Pressey et al. 2007). As a result, species may lose protection as their ranges shift out of current reserve boundaries (Schloss et al. 2011). Therefore the development of complementary efforts that helps decision-makers to identify best areas for restoration activities under a climate change perspective should be taken as a major objective.

The increasing development of freely available spatial software, and the production and release of geospatial data by governments and international organizations have greatly improved our capacity to address spatial conservation planning challenges (Baldwin et al. 2014). Furthermore, the availability of niche modeling softwares and the development and release of future climate projections have also increased our capability for conservation planning under a climate change scenario (Schwartz 2012). Even though, available planning software, such as Marxan are already capable to handle future environmental variability (*e.g.* Carvalho et al. 2011; Veloz et al. 2013), they often need large amount of specific data that may not be readily available, or even may not be relevant for regional scale restoration prioritization goals. Moreover, their use may be perceived as complex and challenging by practitioners, which could preclude their application for decision-making (Baldwin et al. 2014).

To overcome these difficulties, the integration of Multi-Criteria Decision Analysis (MCDA) in a Geographic Information System (GIS) could provide an inexpensive, simple, flexible, transparent and replicable approach to integrate available and generated spatial information to prioritize sites for restoration initiatives at a regional scale. From a rudimentary perspective a GIS-based MCDA or Spatial Multi-Criteria Decision Analysis (SMCDA) can be seen as a decision support process that integrates geospatial data and combining rules to obtain information for decision-making (Malczewski 2006). The SMCDA approach provides a framework that takes explicit account of multiple criteria, helps to structure the management problem, provides a model that can serve as a focus for discussion, and offers a transparent process that leads to rational, justifiable, and explainable decisions (Mendoza & Martins 2006). SMCDA has been increasingly used in environmental sciences and forest management in last decades; however it application for selecting sites for restoration have been scarcely explored (Mendoza & Martins 2006; Huang et al. 2011).

The objective of this study was to develop and evaluate a complementary approach to systematic conservation planning that could be widely used to identify and prioritize site for species restoration initiatives at the regional scale. As a representative case study, we focused our analysis on the selection of priority areas for restoration for two threatened endemic tree species *Bielschmiedia miersii* and *Pouteria splendens* of the “Chilean Rainfall-Valdivian Forest Biodiversity Hotspot” (Arroyo et al. 2004). These species are dominant trees in their respective ecological communities, are inadequately covered by protected areas, and are increasingly threatened by human driven activities (Hechenleitner et al. 2005; Schulz et al. 2010; Pliscoff & Fuentes-Castillo 2011).

## Methodology

### Study area

The study area covers the “shrub and sclerophyllous forest ecological region” of Central Chile (Gajardo 1994), including the coastal and inner territories that make up the current distribution of *B. miersii* and *P. splendens* (Fig. 1). Original landscapes within this ecological region were characterized by a dominance of shrubs species in the coastal ranges, and a mix of forest and shrubs in more inland areas (Gajardo 1994). This region has been severely transformed by the fragmentation of the original landscape, and the remaining habitats are increasingly threatened by human activities (Pliscoff & Fuentes-Castillo 2011).

**Figure 1.**
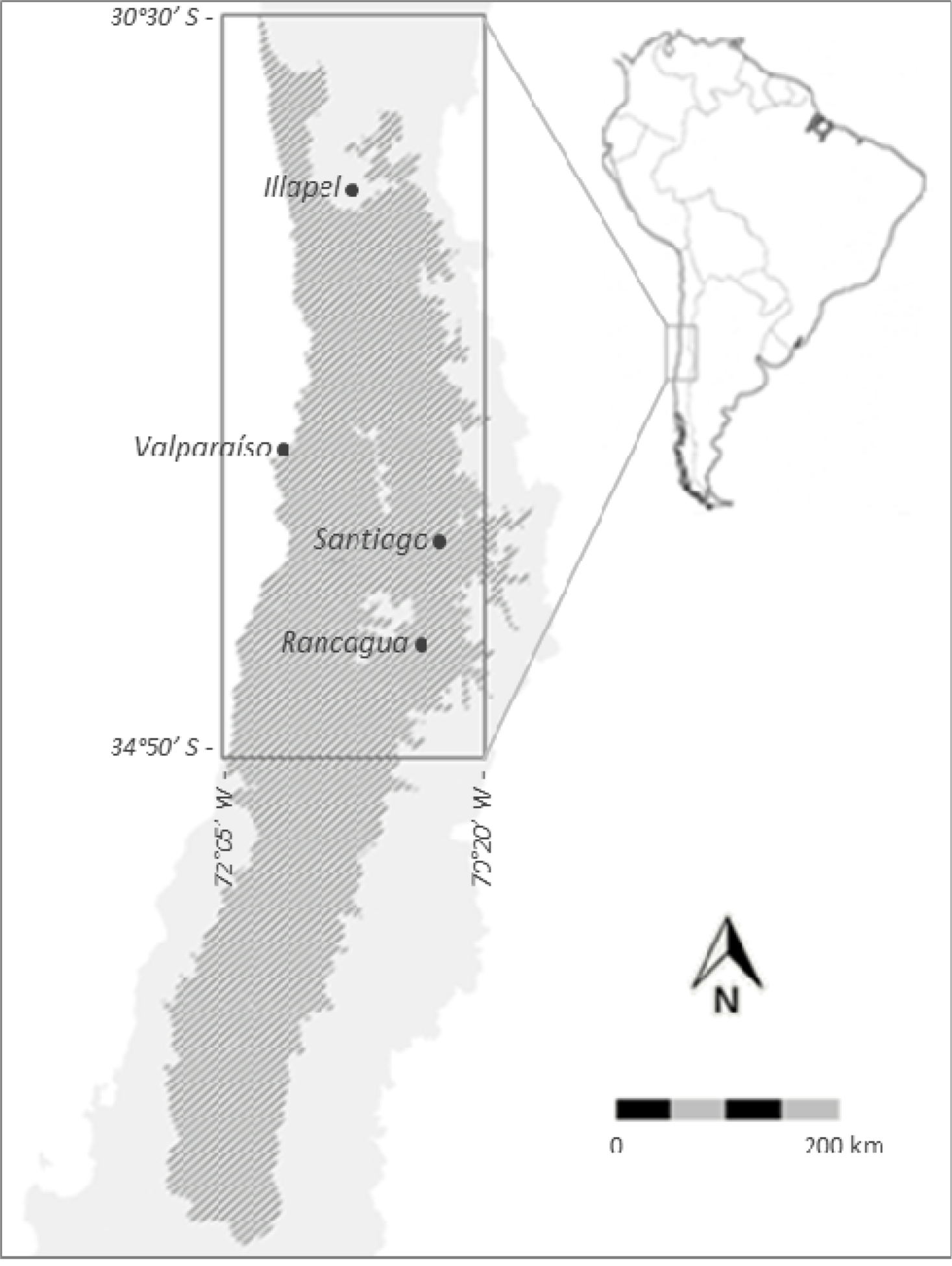
Study Area. The shaded area corresponds to the “shrub and sclerophyllous forest” ecological region, which was used as the boundaries for the niche modeling process. Rectangular area includes the range of distribution of both species, and corresponds to the extent used for creating the maps showed in following sections. Cities mentioned in the text are shown.

Climate within the study area can be broadly characterized as Mediterranean, with marked colder temperatures and rainy periods during winter months, and warmer and dry period during summer (Luebert & Pliscoff 2006). However, local climate characteristics differ considerably over the geographical range of the study area. Historical data for the northern city of Illapel (31°37’50’’ S; 71°09’55’’ W) registers an annual total precipitation of ∼240 mL, while data for the southern city of Rancagua (31°54’43’’ S; 71°30’39’’ W) reports a yearly total precipitation of ∼500 mL. Annual average temperatures are similar over the study range (around 12 - 13°C), but thermic oscillation during warmer and colder seasons differs between the coastal and inland territories (Dirección Meteorológica de Chile 2001). For example, the inner city of Santiago (33°27’50’’ S; 70°38’26’’ W) has an average maximum temperature of 29.7°C during summer months and a minimum average temperature of 3.9° C during winter, whereas for the same period the maximum and minimum temperatures for the coastal city of Valparaiso (33°02’42’’ S; 71°37’14’’ W) are 20.8°C and 9.2°C respectively (Cruz & Calderón 2008).

### Spatial Multi Criteria Decision Analysis (SMCDA)

To identify the best suitable areas for restoration with *B. miersii* and *P. splendens* we performed a GIS-based multi criteria decision analysis (Malczewski 2006) consisting in the integration of four spatial raster layers by using a weighted scheme (Eq. 1).

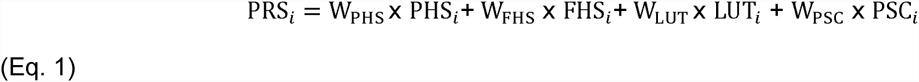

We designed the equation and layers to produce a Priority for Restoration Score (PRS) ranging from 0 to 10 for each species (*i*), with higher values indicating better areas for restoration. Weights (W_*x*_) are specific for each variable and their sum must be equal to 1. Spatial layers in Equation 1 are: Present habitat suitability (PHS), used as an indicator of the relative quality of each pixel’s current climatic conditions for the occurrence of the assessed species. Future habitat suitability (FHS), used as an indicator of the relative quality of each pixel’s future climatic conditions for the occurrence of the assessed species. Land-use type (LUT), which represents the current availability of lands with potential conditions to start a restoration project with the studied species. Priority sites for conservation (PSC), which are areas prioritized by the Chilean government for future conservation initiatives, and used as an indicator of the suitability of each pixel to hold restoration activities in the long-term. The specific methods to generate these four layers are explained below.

#### Present Habitat Suitability (PHS)

We used Maxent software version 3.3.3k (Phillips et al. 2006; Phillips & Dudík 2008; http://www.cs.princeton.edu/∼schapire/maxent) to model present and future potential habitat suitability of *B. miesrsii* and *P. splendens*. We input georeferenced data of recorded individuals of both species as presence points and climatic geospatial data gathered from the WorldClim database (http://www.worldclim.org). Worldclim database consists in 19 bioclimatic layers generated from the interpolation of climatic data compiled around the world with a resolution of ∼1 Km^2^ (Hijmans et al. 2005). Species-presence data was obtained from herbarium specimens, literature records, previous species surveys, and data collected in several field campaigns during the year 2011. We aggregated species presence points to match climatic data resolution avoiding pseudo replication. A total of 75 presence points for *B. miersii* and 22 for *P. splendens* were included in the modeling procedure.

To ensure the quality of the final habitat suitability models and to reduce potential over-parameterization (Williams et al. 2003; Merow et al. 2013) we performed a Pearson correlation analysis of the 19 bioclimatic variables using for that the software ENMTools version 1.4.4 (Warren et al. 2010). As suggested by previous studies (Kumar & Stohlgren 2009), all variables with correlations larger than 0.8 were evaluated to retain only those more relevant for the species ecology. After the correlation analysis, only 8 out the 19 bioclimatic variables were included in the modeling of *B. miersii* and *P. splendens* suitable habitats (See supplementary data).

To select the model parameters we generated and compared several different models by utilizing the corrected Akaike information criterion (AICc) available in the software ENMTOOLS version 1.4.4 (Warren et al. 2010). We took this approach, instead of using Maxent default parameters, as recent studies have shown that could provide better results than Maxent standard configuration (*e.g.* Merow et al. 2013; Syfert et al. 2013), especially when modeling with a small number of samples (< 20 - 25) (Shcheglovitova & Anderson 2013). The results suggested that the best combination of parameters were HQPT with a regularization multiplier of 2 for *B. miersii* and LQ with a regularization multiplier of 1 for *P. splendens* (See supplementary data).

To generate the final model for *B. miersii* we used a random sample of 75% of the presence points, while the 25% remaining points were used to validate the model. The model efficiency was analyzed by the reported AUC (area under the curve). The generated distribution model for *B. miersii* showed an AUC of 0.974, which is classified as an excellent predicting performance (Elith 2000).

In the case of *P. splendens* we followed the “Jackknife” model testing procedure proposed by Pearson et al. (2007), which is especially helpful for bioclimatic modeling with small presence point data. The procedure consists in removing one presence point from the original dataset (n=22) to subsequently run the model with the remainder (n – 1, 21 points) presence points. This process was repeated for all presence points generating 22 different models. We then analyzed *P. splendens* modeling performance by using the software “pValuecompute v.1.0” (Pearson et al. 2007). Performance test for the modeled distribution showed good predicting capability, with a success of 86% (p < 0.001).

Finally we built the PHS raster layers by standardizing the data produced by the Maxent modeling phase into values ranging from 0 to 10. We did this by dividing the probability values of each pixel by the maximum probability value obtained in the entire grid, and then multiplying the resulting value by 10. Generated layer was then resampled to 100 m/pixel through the bilinear interpolation method.

#### Future Habitat Suitability (FHS)

To generate the future habitat suitability layer under a hypothetical climate change scenario, we re-projected the models generated in the PHS section by using projected climatic data for the period 2041- 2060. We used the climatic projection model known as HadGEM2-ES (Jones et al. 2011) because of its good overall performance in predicting the seasonal variability of precipitation and temperature in South America (Cavalcanti & Shimizu 2012). We chose the RCP 2.6 scenario as a conservative representation of concentrations pathways (RCP) of greenhouse gases (Moss et al. 2010). The RCP 2.6 scenario assumes that the global greenhouse emissions will have their maximum concentration between the years (2010 - 2020), declining after this period. The new climatic layers were downscaled and calibrated using WorldClim 1.4 as the baseline “present” climate (Hijmans et al. 2005). The same bioclimatic variables selected to build the distribution model under the current climatic conditions for each species were used to perform the modeling of future potential habitat suitability. The final FSH raster layer was standardized in values ranging between 0 and 10 and resampled to 100 m/pixel by using the same process used to generate the PSH layer.

#### Land-use type (LUT)

We used the Chilean national forest inventory spatial dataset (available at http://sit.conaf.cl/) to categorize land-use within the study area. This polygon-based data set was recently updated and is one of the main tools used by the government to develop policies regarding to forest conservation and management (CONAF-Corporación Nacional Forestal 2011). We reclassified the more than 50 land-use type categories present in the inventory in three main classes (*i.e.* urban, productive, natural) based on the urban-rural-natural spatial pattern present in central Chile (Schulz et al. 2010). Urban lands encompassed all areas classified as urban or industrial by the forest inventory, and were considered not suitable for restoration and therefore given a value of 0. We based this decision on the large difficulties involved in recovering urban and industrial land into areas suitable for restoration initiatives (Pavao-Zuckerman 2008). Productive lands included all areas used for agriculture and silvicultural activities and were given a value of 5. This intermediate score represented areas that could be suitable for restoration, but where restoration projects will face several difficulties due to private land-use conflicts and soil restoration challenges (Rey Benayas & Bullock 2012). Natural lands grouped all areas covered by natural vegetation communities, and were given a value of 10. The resulting reclassified vector layer was then converted to a raster layer with a 100 m/pixel resolution.

#### Priority Sites for Conservation (PSC)

Priority Sites for Conservation are specific areas identified by the Chilean government as the most important zones to develop private or public conservation efforts. There are a total of 68 Priority Sites for Conservation in Chile, and they represent a proactive mechanism specified in the Chilean Biodiversity National Strategy to promote initiatives focused on conservation and protection of biodiversity in the long-term (Conama 2003; Pliscoff & Fuentes-Castillo 2011)We used the latest version of the PSC vector layer available from the Chilean Environmental Ministry map service (http://ide.mma.gob.cl). From this layer we selected only the PSCs that were within our study area. We assigned a value of 10 to all the areas within defined priority sites for conservation, and a decreasing values of one unit (*i.e.* 9,8,7…0) for every one kilometer of distance to the priority area. Therefore, all the areas that are farther than 9 km from a PSC have a value of 0. Our scoring scheme for the PSC layer (*i.e.* distance based) follows the recommendation of the Convention on Biological Diversity (Article 8) to develop complementary sustainable development efforts near protected and other areas of conservation importance (United Nations 1992). The resulting vector layer was then converted to a raster layer with a 100 m/pixel resolution.

### SMCDA Weighting and Analysis

We developed five weighting scenarios to evaluate the effect of different approaches on habitat prioritization results. The weighting scenarios covered a gradient from an extreme short-term to an extreme long-term approach (Table 1). The extreme short-term scenario (EST) allocates all the weight to the layers related with the current feasibility to start restoration projects (*i.e.* PHS, LUT), disregarding the contribution of the layers implicated with the long-term viability of restoration projects. At the other hand, the extreme long-term scenario (ELT) allocates all the weight to the layers associated with long-term viability of restoration activities (*i.e.* FHS, PSC), but disregards factors that may be important for the current implementation of the project. Between these extreme scenarios we built three intermediate weighting scenarios, including a short-term (ST), a non-weighted (NW), and a long-term (LT) scenario (Table 1).

**Table 1.** Scenario weighting scheme used for both species. Scenarios are: EST; extreme short-term, ST; short-term, NW; non-weighted, LT; long-term, ELT; extreme long-term.

| Layers | Scenario Weighting |  |  |  |  |
| --- | --- | --- | --- | --- | --- |
|  | EST | ST | NW | LT | ELT |
| Present Habitat Suitability | 0.50 | 0.40 | 0.25 | 0.20 | 0.00 |
| Future Habitat Suitability | 0.00 | 0.20 | 0.25 | 0.40 | 0.50 |
| Land-Use Type | 0.50 | 0.30 | 0.25 | 0.10 | 0.00 |
| Priority Sites for Conservation | 0.00 | 0.10 | 0.25 | 0.30 | 0.50 |

All GIS processing was performed using the free GIS platform Quantum GIS 2.6 Brighton (www.qgis.org). Output layers generated from the SMCDA were “masked” to fit only the areas potentially suitable for the assessed species. We did this by creating a masking layer composed by the aggregated area of present and future niche modeling distribution. We used the “minimum presence threshold” value to set the distribution boundary for each species. All final raster layers were translated to prioritization maps for qualitative evaluation, and areas corresponding to the highest suitability scores (*i.e.* 9 and 10) where computed for quantitative analysis.

## Results

### Input layer results

The six raster layers we generated to be used as inputs for the SMCDA are shown in Fig. 2. Even though the main objective of this work was to develop a simple method for prioritizing site for restoration under different scenarios, we consider it relevant to briefly describe the resulting layers that were used as the input variables for our method. This is not only important for placing results in context, but also to provide information that helps to evaluate the SMCDA results.

**Figure 2.**
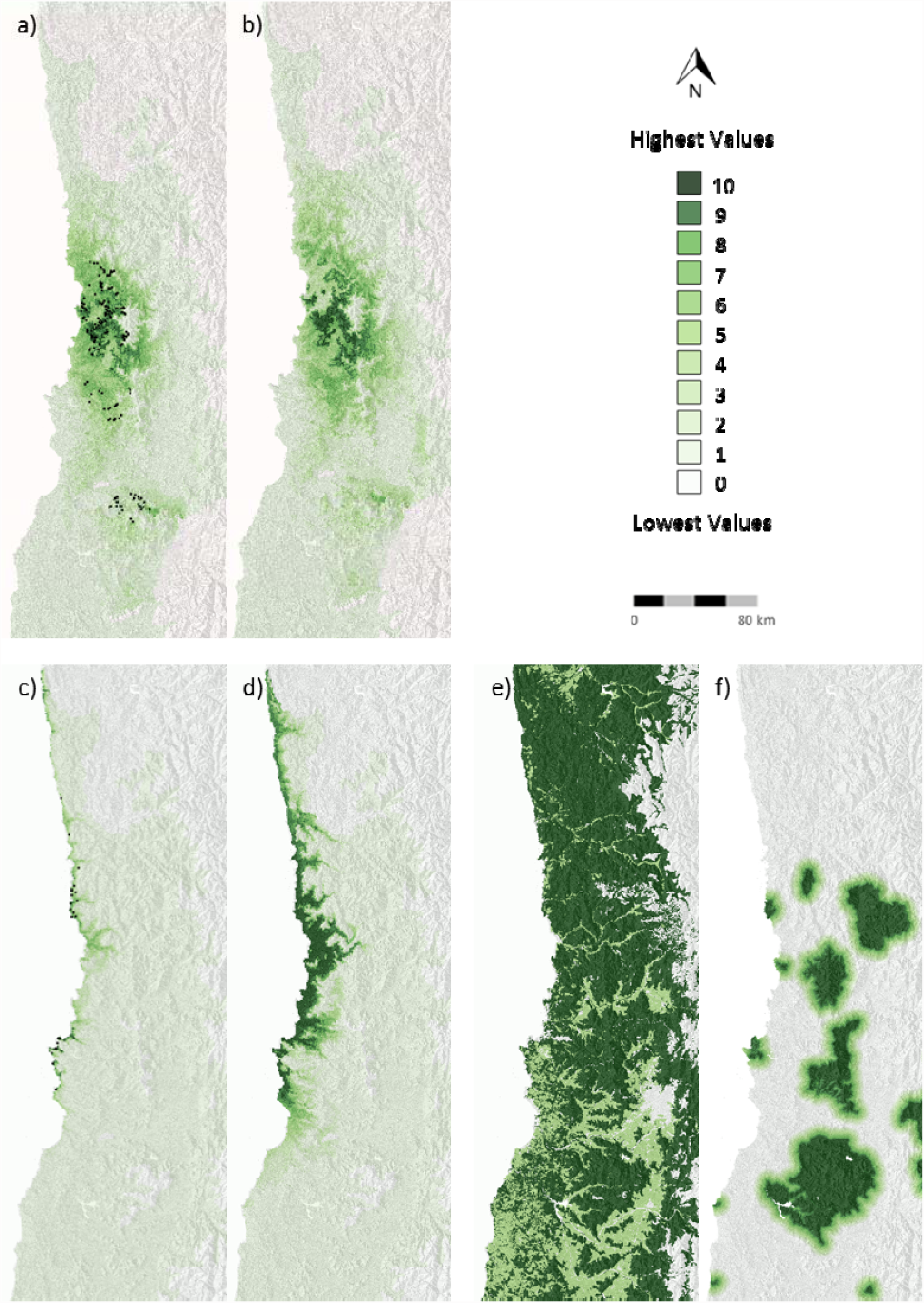
Visualization of the spatial layers built and used as input for the SMCDA. Letters a) and b) represent the modeled present (PHS) and future (FHS) habitat suitability for *B. miersii.* Letters c) and d) represent the present and future (PHS and FHS) modeled habitat suitability for *P. splendens.* In a) and c) black dots correspond to the record presence points of these species that were used for the niche modeling process. Letter e) corresponds to the land-use type (LUT) layer, and letter f) to the priority sites for conservation (PSC) layer.

Modeled distribution under current climate of *B. miersii* shows a concentration of highest values in the mountainous range covering the northwestern part of the species current distribution. This is coincident with the area that concentrates the large extent of recorded populations. However, the model also assigned intermediate and lower values to several areas were currently populations occur, as those in the central and southern part of current distribution (Fig. 2a). The projected distribution with climate change does not show major differences with the projection under current climatic conditions, except for a slight increase in the range of highest values toward the south and east (Fig. 2b).

In the case of *P. splendens*, the modeled distribution under current climate is almost entirely restricted to the coastal plains, ravines, and valleys that are directly influenced by the ocean climate. Highest values for *P. splendens* tend to be localized in three main areas located in the northern, central and southern range of predicted distribution. Whereas the southern and central areas coincide with historic presence records for the species, there are no historic records for the northern area (Fig. 2c). In contrast with *B. miersii*, the modeled distribution for *P. splendens* under climate change shows important changes when compared to the current climate conditions, experiencing a general increase of highest values towards the east, which are mostly concentrated in the central part of the species current distribution range (Fig. 2d).

Land-use types presenting the highest values tend to be concentrated in the northern part of the study area. These high value areas present a gradual reduction of prevalence towards southern zones where seems to be confined mostly to the upper part of valleys (Fig. 2e). Areas presenting land-use types with intermediate values are mostly represented by valleys and flat areas which are majorly concentrated in the southern part of the study area. Areas presenting the lowest values are concentrated in the eastern part of the area of study. This is coincident with the presence of the Andes Mountain Range, which due to their altitude, topography and severe climatic conditions generate different land covers (e.g. volcanic debris, glaciers, bare rock) that are not viable for restoration with the assessed species (Fig. 2e).

Areas defined as priority sites for conservation (PSC) by the Chilean government are not evenly distributed in the study area, which is clearly seen as the concentration of highest values in the central and southern parts of the study area, and a complete absence of these sites in northern area (Fig. 2f). Furthermore, PSC are mostly concentrated in the coastal mountainous range (between the Andes mountain and the coast), whereas the coastal plains and zones adjacent to the ocean present only small and highly isolated PSC (Fig. 2f).

### SMCDA results

The generated Priority for Restoration Scores (PRS) maps for the five weighting schemes for *B. miersii* and *P. splendens* are shown in Figure 3. These maps show the distribution of PRS, which represent the ranked suitability of different areas for developing restoration activities under the five different weighting scenarios.

**Figure 3.**
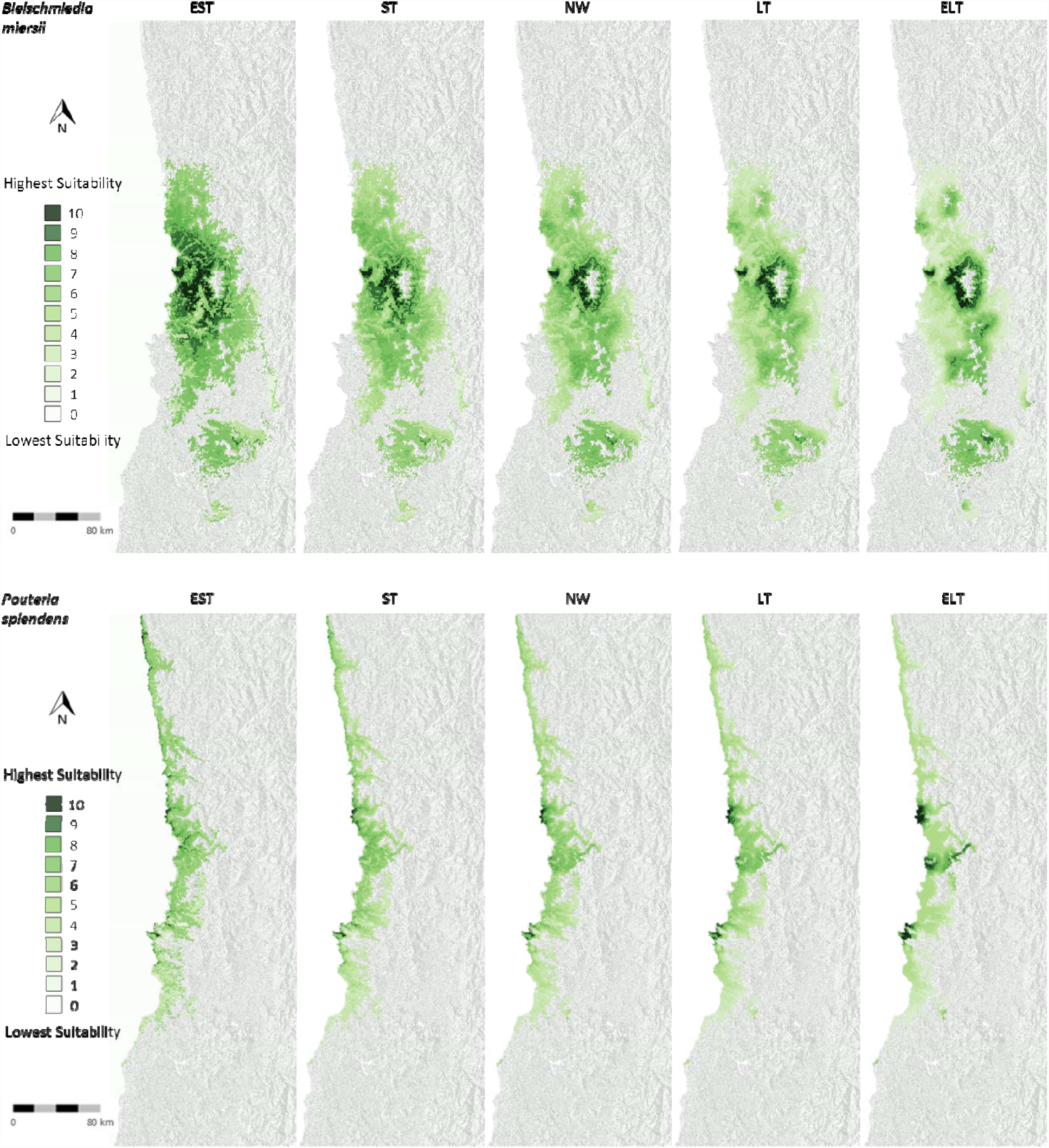
Suitability of areas for restoration with *B. miersii* (top) and *P. splendens* (bottom) under the five assessed scenarios generated through the SMCDA. Values represent Priority for Restoration Score (PRS). Scenarios are: EST; extreme short-term, ST; short-term, NW; non-weighted, LT; long-term, ELT; extreme long-term.

There are important differences in the spatial patterns of PRS between the different weighting scenarios. In general, both for *B miersii* and *P. splendens*, there is a decreasing average pixel PRS and increasing spatial clustering when moving from the EST to the ELT scenario (Table 2). The fragmented patterns of highest suitable areas shown in short-term scenarios are associated with the projected spatial distribution of current population (Figure 2a,c), whereas the clustered pattern of long-term scenarios are highly associated with the presence of priority sites for conservation (Figure 2,f).

**Table 2.** Main statistical summary of PRS for the five evaluated scenarios for each of the assessed species. Mean, coefficient of variation, and Moran’s I coefficient of autocorrelation are shown. Scenarios are: EST; extreme short-term, ST; short-term, NW; non-weighted, LT; long-term, ELT; extreme long-term.

|  | *B. miersii* |  |  |  |  | *P. splendens* |  |  |  |  |
| --- | --- | --- | --- | --- | --- | --- | --- | --- | --- | --- |
|  | EST | ST | NW | LT | ELT | EST | ST | NW | LT | ELT |
| Mean | 6.19 | 5.35 | 5.15 | 4.50 | 4.09 | 4.73 | 4.42 | 4.31 | 4.07 | 3.88 |
| CV | 0.29 | 0.31 | 0.33 | 0.41 | 0.58 | 0.37 | 0.35 | 0.35 | 0.41 | 0.51 |
| Moran's I | 0.93 | 0.96 | 0.97 | 0.98 | 0.99 | 0.88 | 0.94 | 0.95 | 0.98 | 0.98 |

While the PRS under the five scenarios show general spatial patterns common to both species, the amount of areas prioritized in the two highest suitability scores does not follow the same patterns (Fig 4). In the case of *B. miersii* the scenario prioritizing the larger amount of areas with the two highest suitability scores (*i.e.* 9 and 10) is the EST, whereas for *P. splendens* is the ELT, which is the opposite scenario. At the other hand, the scenario prioritizing the smaller amount of areas for *B. miersii* is the LT, whereas for *P. splendens* is the ST.

**Figure 4.**
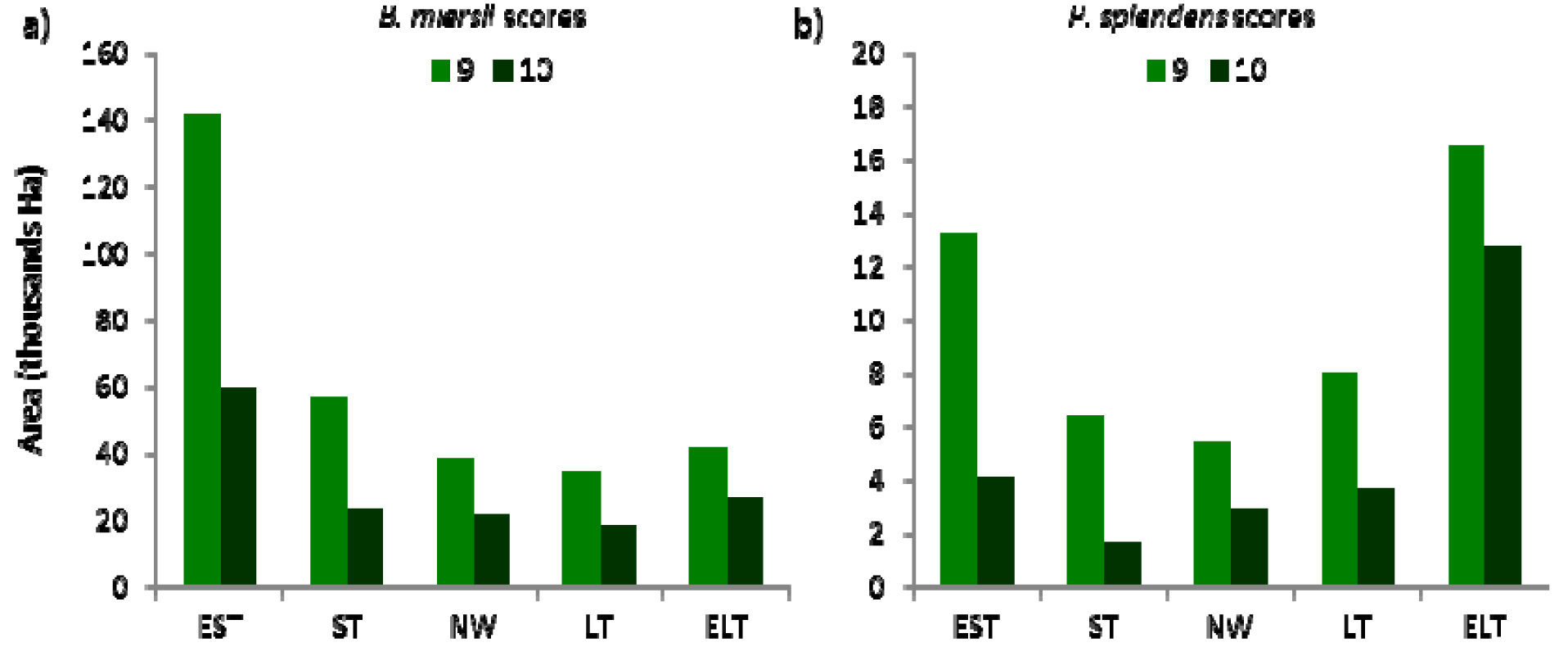
Quantification of total area categorized in the two highest PRS scores (9 and 10) under the five scenarios for each of the two assessed species, a) *B. miersii*, b) *P. splendens.*

## Discussion

Results from our work highlight the usefulness of using a SMCDA approach to identify, evaluate and prioritize sites for restoration at a regional scale. By using this approach we were able to identify and quantify the best suitable areas for restoration initiatives of two threatened endemic species of the Chilean Biodiversity Hotspot. The SMCDA provided a simple and transparent methodological framework to integrate available spatial information, generating insightful knowledge readily usable by local decision-makers. Although we could have use additional spatial information as input layers for our approach, we attempted to focus our analysis to spatial layers that can be gathered or easily generated elsewhere. In view of that the aim of this work was not to present the proposed method as a definitive approach – and neither our maps as definitive results, but rather to put in perspective the usefulness of the SMCDA as a tool for spatial prioritization of sites for biodiversity restoration.

One of the key steps in the developing of a multi-criteria spatial analysis for conservation planning should be the selection of the input layers and the specific weighing applied to each of them (Phua & Minowa 2005; Huang et al. 2011). These decisions do not only have to be focused on combining relevant available information, but also must produce legitimate results for decision makers (Munda 2005).

Therefore, the relevancy, accuracy and reliability of the information contained in the input layers are key factors for the quality and credibility of our results

### Input layers

While we could theoretically include a large number of input variables in our SMCDA, the limited availability of spatial information with adequate resolution importantly reduced the number of potential input layers we could use. Furthermore, because the scope of our work was to develop a methodological framework that can be used elsewhere, selection of input layers not only had to be based in the available spatial information in our study area, but also in other regions with similar conservation challenges.

In this regard, we are aware that several regions may not have updated and accurate available land-use layers as we had, or if they have, the accessibility to the information could be restricted to governmental agencies. However this limitation could be solved by using freely available land-classification software and satellite images, which is a methodology that can accurately classify land cover in the three categories we used in our approach (*e.g.* Nolè et al. 2015). Even though the of land-use in only three categories (i.e. urban, rural, natural) can be considered too broad for local-scale conservation planning, it could provide useful information about regional-scale availability of natural lands that can be currently targeted for restoration activities with focal species.

The inclusion in the SMCDA of official recognized “priority sites for conservation (PSC)” is one of the major novelties of our approach. In contrast with traditional protected areas, PSC are mostly areas not currently protected, but officially recognized as primary importance to be protected in the short to mid-term (Conama 2003; Tognelli et al. 2008). PSC inclusion provide an objective indicator on the specific areas that governments would support for future conservation initiatives, which is a fundamental information for planning restoration initiatives under future climate change scenarios. Because the designation of PSC is based in the Convention on Biological Diversity (United Nations 1992; article 8 letters a and b), PSC’s represent information that should be available in several of the 194 countries signatories of this convention, and therefore supports its inclusion in the SMCDA as a commonly available spatial information.

The use of niche modeling software is a key part of our approach because provides fundamental ecological information regarding the areas that have current and future environmental conditions to potentially support restoration projects with the focal species. Additionally, results from the niche modeling phase, such as the most relevant environmental variables related to each species, may also contribute to understand what are key environmental conditions related with the occurrence of species, which can provide useful ecological information for the development of local scale restoration management strategies (Schwartz 2012). However, while niche modelling software are powerful tools, the accuracy of predicted distribution results will depend on the quality of presence data, environmental layers, and parameter settings (Loiselle et al. 2008; Costa et al. 2010; Soria-Auza et al. 2010) In this regard, researchers and practitioners need to be particularly cautious of potential data flaws if they are to use niche modelling to generate the species distribution layers to be used in the SMCDA.

### Scenarios weighting scheme

The main characteristic of our weighting scheme was to explicitly relate the weighting scenarios with the underlying characteristics of the used spatial layers in terms of their relevance for short-term and long-term decision-making. Each of the five built scenarios has an implicit narrative behind that helps to explain decision-makers the specific weight assigned to each variable. Our assumption was that if restoration strategies are based in short-term planning, current environmental conditions and available natural lands will prevail in decisions. However, if strategies are developed to embrace long-term perspectives, future environmental conditions and projected protected lands need to be increasingly taken into account.

Even though this weighting scheme was designed to be easily communicated to decision-makers, we did not include a participatory phase to integrate potential decision-makers diversity of opinions. This participatory phase is often considered a fundamental step to legitimate the results in building conservation policies (Munda 2005; Ananda & Herath 2008). However, as we have stated before, our work was not focused on generating a definitive outcome, but rather on evaluating the potential application of the SMCDA approach as a tool for restoration prioritization initiatives. Although our weighting scheme has not passed through a legitimatization phase by decision-makers, we still consider that our results could be helpful because the gradient weighting scheme provide decision-makers with a range of outcomes that can be used as exploratory boundaries to evaluate potential results. Indeed participatory phases often derive in a range of weights, and thus in more than one set of results that are finally used to make a decision (Chen et al. 2010). The use of a gradient weighting scheme when using a small number of variables could also be used to evaluate the sensibility of the SMCDA to changes in weighting values, and therefore to understand what specific values play larger roles in generated results.

### SMCDA results

Outcomes generated through the SMCDA reveals the specific effects of weighting scenarios and the spatial interaction of spatial layers in the distribution and extent of areas identified with high priorities for developing restorations initiatives with the two species used in our study. These results not only provide useful information for local decision-makers, but also help to understand the role of the different input layers in final outcomes.

Results of the SMCDA for *B. miersii* show a significant larger amount of prioritized area for restoration when compared with the suitable areas for *P. splendens* independently of the assessed scenario, which was an expected result based in the much larger distribution of the former species compared to the later. However, an interesting result from our work was that the total suitability areas for restoration of each species changed in opposite direction when moving from the short-term to the long-term weighting scenarios. Whereas weighting emphasizing long-term scenarios tended to reduce the availability of suitable habitats for *B. miersii* restoration, these same scenarios increase the extent of suitable areas for *P. splendens* restoration projects. As two of out of the four layers (*i.e.* PSC, LUT) were shared for both species in the SMCDA, the divergences of these results are mainly related with the change on the species distribution due to the climate change scenario. In fact, while *B. miersii* is predicted to see reductions in its potential distribution under the future climate, *P. splendens* is expected to have an opposite response, presenting a large increase in projected distribution. This species specific response to climate change highlights the importance to take into account the future climate variability for planning plant species restoration initiatives (Gelviz-Gelvez et al. 2015). Because these two species are dominant trees in their respective ecological community, they can be used as proxy for selecting priority sites for restoration efforts focused in the entire vegetation community.

## Conclusion

In this work we demonstrate the usefulness of integrating available spatial layers and niche modeling into a SMCDA approach to develop an affordable, flexible, transparent, and replicable method to prioritize areas for restoration initiatives of plant species taking into account the future climatic variability. It is affordable because it can be performed by using free software (*i.e.* Maxent, QGIS) and freely available spatial information. It is flexible because input layers, layer scoring, and weighting schemes can be modified to fit specific decision-making contexts. It is transparent because each of the steps is clearly identified and justified. And it is replicable because it uses information that can be gathered or generated elsewhere. Finally, our approach does not aim to be used in replacement of other local-scale software-based planning approach, such as Zonation and Marxan, but rather as a complementary method that bridge the global-scale priority areas, with local-scale restoration planning efforts. This SMCDA approach could be used as a management tool to prioritize regional-scale areas for restoration of plant species in Chile, as well as in other countries with restricted budgets for conservation efforts.

## Acknowledgments

We would like to thank Gloria Montenegro, Eduardo Arellano and Luis Olivares from the Faculty of Agronomy and Forestry Engineering of the Pontifical Catholic University of Chile for their support during this research. This project was partially funded by the Chilean Native Forest Research Grant, Project CONAF 025/2010: “Distribución, hábitat potencial y diversidad genética de poblaciones de Belloto del Norte (Beilschmiedia miersii) y Lúcumo chileno (Pouteria splendens)”,

## References

Ananda J, Herath G (2008) Multi-attribute preference modelling and regional land-use planning. Ecol Econ 65:325–335. doi: 10.1016/j.ecolecon.2007.06.024

Arroyo MTK, Marquet PA, Marticorena C, et al. (2004) Chilean winter rainfall-valdivian forests. Pages 99–103 In: Mittermeier RA, Gil PR, Hoffmann M, et al. (eds) Hotspots Rev. Earth’s Biol. Wealthiest Most Threat. Ecosyst. Sierra-Cemex, Mexico.

Baldwin R, Scherzinger R, Lipscomb D, et al. (2014) Planning for Land Use and ConservationL: Assessing GIS-Based Conservation Software for Land Use Planning. Fort Collins, CO

Brook BW, Sodhi NS, Bradshaw CJA (2008) Synergies among extinction drivers under global change. Trends Ecol Evol 23:453–60. doi: 10.1016/j.tree.2008.03.011

Brooks TM, Mittermeier RA, da Fonseca GAB, et al. (2006) Global Biodiversity Conservation Priorities. Science (80-) 313:58–61. doi: 10.1126/science.1127609

Carvalho SB, Brito JC, Crespo EG, et al. (2011) Conservation planning under climate change: Toward accounting for uncertainty in predicted species distributions to increase confidence in conservation investments in space and time. Biol Conserv 144:2020–2030. doi: 10.1016/j.biocon.2011.04.024

Cavalcanti IF, Shimizu MH (2012) Climate Fields over South America and Variability of SACZ and PSA in HadGEM2-ES. Am J Clim Chang 01:132–144. doi: 10.4236/ajcc.2012.13011

Chen Y, Yu J, Khan S (2010) Spatial sensitivity analysis of multi-criteria weights in GIS-based land suitability evaluation. Environ Model Softw 25:1582–1591. doi: 10.1016/j.envsoft.2010.06.001

CONAF-Corporación Nacional Forestal (2011) Catastro de los Recursos Vegetacionales Nativos de Chile. Gobierno de Chile, Santiago, Chile

Conama (2003) National Biodiversity Strategy of the Republic of Chile.

Gobierno de Chile, Santiago, Chile Costa GC, Nogueira C, Machado RB, Colli GR (2010) Sampling bias and the use of ecological niche modeling in conservation planning: A field evaluation in a biodiversity hotspot. Biodivers Conserv 19:883–899. doi: 10.1007/s10531-009-9746-8

Cruz C, Calderón J (2008) Guía Climatológica Práctica. Gobierno de Chile, Santiago, Chile

DMC-Dirección Meteorológica de Chile (2001) Climatología Regional. Gobierno de Chile, Santiago, Chile

Elith J (2000) Quantitative methods for modeling species habitat: Comparative performance and an application to Australian plants. Pages 39–58 In: Ferson S, Burgman M (eds) Quant. Methods Conserv. Biol. Springer, New York

Ellis EC, Klein Goldewijk K, Siebert S, et al. (2010) Anthropogenic transformation of the biomes, 1700 to 2000. Glob Ecol Biogeogr 19:589–606. doi: 10.1111/j.1466-8238.2010.00540.x

Ellis EC, Ramankutty N (2008) Putting people in the map: anthropogenic biomes of the world. Front Ecol Environ 6:439–447. doi: 10.1890/070062

Foley JA, Defries R, Asner GP, et al. (2005) Global consequences of land use. Science (80-) 309:570–574. doi: 10.1126/science.1111772

Gajardo R (1994) La vegetación natural de Chile: clasificación y distribución geográfica. Editorial Universitaria, Santiago, Chile

Gelviz-Gelvez SM, Pavón NP, Illoldi-Rangel P, Ballesteros-Barrera C (2015) Ecological niche modeling under climate change to select shrubs for ecological restoration in Central Mexico. Ecol Eng 74:302–309. doi: 10.1016/j.ecoleng.2014.09.082

Hechenleitner P, Gardner M, Thomas P, et al. (2005) Plantas Amenazadas del Centro-sur de Chile. Universidad Austral de Chile, Valdivia, Chile

Hijmans RJ, Cameron SE, Parra JL, et al. (2005) Very high resolution interpolated climate surfaces for global land areas. Int J Climatol 25:1965–1978. doi: 10.1002/joc.1276

Hoekstra JM, Boucher TM, Ricketts TH, Roberts C (2005) Confronting a biome crisis: global disparities of habitat loss and protection. Ecol Lett 8:23–29. doi: 10.1111/j.1461-0248.2004.00686.x

Holl K, Crone E, Schultz C (2003) Landscape restoration: moving from generalities to methodologies. BioScience 53:491–502.

Huang IB, Keisler J, Linkov I (2011) Multi-criteria decision analysis in environmental sciences: Ten years of applications and trends. Sci Total Environ 409:3578–3594. doi: 10.1016/j.scitotenv.2011.06.022

Jones CD, Hughes JK, Bellouin N, et al. (2011) The HadGEM2-ES implementation of CMIP5 centennial simulations. Geosci Model Dev Discuss 4:689–763. doi: 10.5194/gmdd-4-689-2011

Kukkala AS, Moilanen A (2013) Core concepts of spatial prioritisation in systematic conservation planning. Biol Rev 88:443–464. doi: 10.1111/brv.12008

Kumar S, Stohlgren TJ (2009) Maxent modeling for predicting suitable habitat for threatened and endangered tree Canacomyrica monticola in New Caledonia. J Ecol Nat Sci Vol 1(4) 1:094–098. doi: 10.3390/d1020118

Loiselle BA, Jørgensen PM, Consiglio T, et al. (2008) Predicting species distributions from herbarium collections: Does climate bias in collection sampling influence model outcomes? J Biogeogr 35:105–116. doi: 10.1111/j.1365-2699.2007.01779.x

Luebert F, Pliscoff P (2006) Sinopsis bioclimática y vegetacional de Chile. Editorial Universitaria, Santiago, Chile Malczewski J (2006) GIS-based multicriteria decision analysis: a survey of the literature. Int J Geogr Inf Sci 20:703–726. doi: 10.1080/13658810600661508

Margules CR, Pressey RL (2000) Systematic conservation planning. Nature 405:243–53. doi: 10.1038/35012251

Mendoza GA, Martins H (2006) Multi-criteria decision analysis in natural resource management: A critical review of methods and new modelling paradigms. For Ecol Manage 230:1–22. doi: 10.1016/j.foreco.2006.03.023

Merow C, Smith MJ, Silander JA (2013) A practical guide to MaxEnt for modeling species’ distributions: What it does, and why inputs and settings matter. Ecography (Cop) 36:1058–1069. doi: 10.1111/j.1600-0587.2013.07872.x

Mittermeier RAA, Gil PR, Hoffman M, et al. (2004) Hotspots RevisitedL: Earth’s biologically richest and most endangered ecoregions. Cemex-Conservation International, México

Moss RH, Edmonds JA, Hibbard KA, et al. (2010) The next generation of scenarios for climate change research and assessment. Nature 463:747–756. doi: 10.1038/nature08823

Munda G (2005) Multiple criteria decision analysis and sustainable development. Mult. criteria Decis. Anal. State art Surv. Springer New York, New York

Myers N (1990) The Biodiversity Challenge: Expanded Hot-Spots Analysis. Environmentalist 10:243–256.

Myers N, Mittermeier RA, Mittermeier CG, et al. (2000) Biodiversity hotspots for conservation priorities. Nature 403:853–858. doi: 10.1038/35002501

Nolè G, Murgante B, Calamita G, et al. (2015) Evaluation of urban sprawl from space using open source technologies. Ecol Inform 26:151–161. doi: 10.1016/j.ecoinf.2014.05.005

Noss R, Nielsen S, Vance-Borland K (2009) Prioritizing ecosystems, species, and sites for restoration. In: Moilanen A, Wilson K, Possingham H (eds) Spat. Conserv. Prioritization Quant. Methods Comput. Tools. Oxford University Press, Oxford, UK, pp 158–171

Olson D, Dinerstein E (2002) The Global 200: Priority ecoregions for global conservation. Ann Missouri Bot Gard 89:199–224.

Pavao-Zuckerman MA (2008) The Nature of Urban Soils and Their Role in Ecological Restoration in Cities. Restor Ecol 16:642–649. doi: 10.1111/j.1526-100X.2008.00486.x

Pearson RG, Raxworthy CJ, Nakamura M, Townsend Peterson A (2007) Predicting species distributions from small numbers of occurrence records: a test case using cryptic geckos in Madagascar. J Biogeogr 34:102–117. doi: 10.1111/j.1365-2699.2006.01594.x

Phillips S, Dudík M (2008) Modeling of species distributions with Maxent: new extensions and a comprehensive evaluation. Ecography (Cop) 161–175. doi: 10.1111/j.2007.0906-7590.05203.x

Phillips SJ, Anderson RP, Schapire RE (2006) Maximum entropy modeling of species geographic distributions. Ecol Modell 190:231–259. doi: 10.1016/j.ecolmodel.2005.03.026

Phua M-H, Minowa M (2005) A GIS-based multi-criteria decision making approach to forest conservation planning at a landscape scale: a case study in the Kinabalu Area, Sabah, Malaysia. Landsc Urban Plan 71:207–222. doi: 10.1016/j.landurbplan.2004.03.004

Pliscoff P, Fuentes-Castillo T (2011) Representativeness of terrestrial ecosystems in Chile’s protected area system. Environ Conserv 38:303–311. doi: 10.1017/S0376892911000208

Pressey RL, Bottrill MC (2009) Approaches to landscape- and seascape-scale conservation planning: convergence, contrasts and challenges. Oryx 43:464. doi:10.1017/S0030605309990500

Pressey RL, Cabeza M, Watts ME, et al. (2007) Conservation planning in a changing world. Trends Ecol Evol 22:583–92. doi:10.1016/j.tree.2007.10.001

Redford KH, Coppolillo P, Sanderson EW, et al. (2003) Mapping the Conservation Landscape. Conserv Biol 17:116–131. doi:10.1046/j.1523-1739.2003.01467.x

Rey Benayas JM, Bullock JM (2012) Restoration of Biodiversity and Ecosystem Services on Agricultural Land. Ecosystems 15:883–899. doi:10.1007/s10021-012-9552-0

Sarkar S, Pressey RL, Faith DP, et al. (2006) Biodiversity Conservation Planning Tools: Present Status and Challenges for the Future. Annu Rev Environ Resour 31:123–159. doi:10.1146/annurev.energy.31.042606.085844

Schloss CA, Lawler JJ, Larson ER, et al. (2011) Systematic conservation planning in the face of climate change: bet-hedging on the Columbia Plateau. PloS one 6:e28788. doi:10.1371/journal.pone.0028788

Schulz JJ, Cayuela L, Echeverria C, et al. (2010) Monitoring land cover change of the dryland forest landscape of Central Chile (1975–2008). Appl Geogr 30:436–447. doi:10.1016/j.apgeog.2009.12.003

Schwartz MW (2012) Using niche models with climate projections to inform conservation management decisions. Biol Conserv 155:149–156. doi:10.1016/j.biocon.2012.06.011

Shcheglovitova M, Anderson RP (2013) Estimating optimal complexity for ecological niche models: A jackknife approach for species with small sample sizes. Ecol Modell 269:9–17. doi:10.1016/j.ecolmodel.2013.08.011

Soria-Auza RW, Kessler M, Bach K, et al. (2010) Impact of the quality of climate models for modelling species occurrences in countries with poor climatic documentation: a case study from Bolivia. Ecol Modell 221:1221–1229. doi:10.1016/j.ecolmodel.2010.01.004

Syfert MM, Smith MJ, Coomes DA (2013) The Effects of Sampling Bias and Model Complexity on the Predictive Performance of MaxEnt Species Distribution Models. PloS one. doi:10.1371/journal.pone.0055158

Tognelli MF, De Arellano PIR, Marquet P a. (2008) How well do the existing and proposed reserve networks represent vertebrate species in Chile? Divers Distrib 14:148–158. doi:10.1111/j.1472-4642.2007.00437.x

United Nations (1992) Convention on Biological Diversity. https://www.cbd.int/convention/text/default.shtml (accessed 15 July 2015)

Veloz SD, Nur N, Salas L, et al. (2013) Modeling climate change impacts on tidal marsh birds: restoration and conservation planning in the face of uncertainty. Ecosphere 4:1–25. doi:10.1890/ES12-00341.1

Waldron A, Mooers AO, Miller DC, et al. (2013) Targeting global conservation funding to limit immediate biodiversity declines. Proc Natl Acad Sci 110:1–5. doi:10.5061/dryad.p69t1

Warren DL, Glor RE, Turelli M (2010) ENMTools: a toolbox for comparative studies of environmental niche models. Ecography (Cop) 1:607–611. doi:10.1111/j.1600-0587.2009.06142.x

Watson J, Grantham H, Wilson K, Possingham H (2011) Systematic conservation planning: past, present and future. In: Ladle R, Whittaker R (eds) Conserv. Biogeogr. Blackwell Publishing Ltd., pp 136–160

Williams SE, Bolitho EE, Fox S (2003) Climate change in Australian tropical rainforests: an impending environmental catastrophe. Proc Biol Sci 270:1887–92. doi:10.1098/rspb.2003.2464

Wilson K a., Carwardine J, Possingham HP (2009) Setting conservation priorities. Ann N Y Acad Sci 1162:237–264. doi:10.1111/j.1749-6632.2009.04149.x

Wilson KA, McBride MF, Bode M, Possingham HP (2006) Prioritizing global conservation efforts. Nature 440:337–340. doi:10.1038/nature04366

